# Cognitive tests of pattern separation as sleep-sensitive measures in middle-aged and older adults: An exploratory study

**DOI:** 10.1101/2024.08.13.607801

**Authors:** Aina E. Roenningen, Devan Gill, Brianne A. Kent

## Abstract

Sleep disturbances are considered both a dementia risk factor and symptom. The present research aimed to identify cognitive tests sensitive to objective sleep quantity or quality, focusing on tests designed to evaluate the earliest cognitive changes in dementia. We recruited adults who were 50 years or older and remotely monitored their sleep patterns for 7 consecutive days using wrist actigraphy and sleep diaries. On day 7, participants completed different cognitive tests; the Psychomotor Vigilance Task (PVT), the Prodromal Alzheimer’s and Mild Cognitive Impairment battery from the Cambridge Neuro-psychological Test Automated Battery (CANTAB), the Mnemonic Similarity Task (MST), taxing pattern separation, and the Montreal Cognitive Assessment (MoCA). The final sample included 34 participants (mean age: 65.56, SD: 9.57). There were significant correlations between objective sleep quantity and PVT and MST performance. MoCA scores were correlated with performance on CANTAB and MST. Objective sleep quantity predicted MST performance when controlling for age, sex, and education. Tests of pattern separation are sensitive to objective sleep quantity and early cognitive changes associated with dementia. The sensitivity of pattern separation tests to sleep should be evaluated in a larger study, aiming to identify suitable clinical trial outcome measures for sleep-promoting treatments in older adults.

**Statement of Significance:** There is growing emphasis on the importance of sleep as a potential therapeutic target for neurodegenerative diseases (e.g., Alzheimer’s disease). Identifying cognitive measures that are sensitive to both sleep and the earliest cognitive changes associated with dementia are needed for use as outcome measures in clinical trials evaluating the effectiveness of sleep-promoting interventions., Our research addresses this need by investigating the relationship between sleep patterns and performance on cognitive tests designed to assess the earliest cognitive changes in dementia. Our results suggest that cognitive tests designed to assess pattern separation are uniquely sensitive to sleep quantity in older adults, and warrant follow-up with a larger study.

## 1. Introduction

Sleep has recently been identified as a potential therapeutic target for dementia caused by Alzheimer’s disease (Kent et al., 2021). There is growing evidence from humans and preclinical mouse models that sleep can directly affect the accumulation of pathological amyloid-beta and tau proteins in the brain, both of which are the defining hallmarks of AD and are thought to drive the neuropathologic course of the disease (Kent et al., 2021). It is estimated that by treating sleep disorders, 15% of AD cases could be delayed in onset or prevented (Bubu et al., 2017). In addition to the direct effects on neuropathology, there is evidence that sleep disruption may be the mechanistic pathway through which AD pathology affects hippocampal-dependent cognitive decline (Mander et al., 2015). There are now a variety of interventions, from non-invasive brain stimulation (e.g., transcranial electrostimulation) to pharmacological agents (e.g., orexin receptor antagonists, melatonin, or trazodone), being evaluated for safety and effectiveness of improving sleep in individuals diagnosed with AD (Eyob et al., 2024; Kent et al., 2021; McCleery & Sharpley, 2020). The ultimate goal of many of these sleep-focused trials is to improve cognition (i.e., prevent, slow, or reverse the dementia caused by AD) (Eyob et al., 2024). What is now needed is to identify which of the cognitive tests would be most suitable as clinical trial outcome measures, such that the tests are sensitive to the cognitive decline associated with AD and also to sleep-associated cognitive processes.

Research on sleep and cognition in older adults suggests that disruptions in sleep, especially short or long sleep duration, higher fragmentation, and sleep-disordered breathing, contribute to the emergence of cognitive decline (reviewed by Dzierzewski et al., 2022; Scullin & Bliwise, 2015). Meta-analyses have found associations between sleep and overall cognitive function (e.g., MMSE), executive functions, verbal memory, and working memory capacity in older adults (reviewed by Lo et al., 2016). For example, the Sleep and Dementia Consortium combined data from 5 independent community-based cohorts of middle-aged to older adults and found that better sleep quality was associated with better global cognition and that short total sleep time was associated with poorer attention and processing speed (Pase et al., 2023).

Pattern separation is a computational mechanism taking place during encoding and proposed to underlie the creation of distinct ensemble neural representations from overlapping inputs (Marr, 1969; O’Reilly & McClelland, 1994). By transforming similar experiences into discrete neural representations, pattern separation is postulated to increase the likelihood of accurate memory encoding and subsequent retrieval, which is fundamental to successful episodic memory.

Cognitive tests designed to assess pattern separation are sensitive to the cognitive changes associated with mild cognitive impairment (MCI) and AD (Ally et al., 2013; Newsome et al., 2012; Yassa et al., 2010). Performance has been shown to correlate with levels of cerebrospinal fluid amyloid-beta 42, which is a clinically validated AD biomarker (Wesnes et al., 2014). Individuals with MCI and AD also have higher rates of false memories (Budson et al., 2001; Gold et al., 2007; Hart et al., 1985; Hildebrandt et al., 2009; Plancher et al., 2009), which is hypothesized to be caused by a failure in pattern separation leading to heightened interference amongst memories (Kent et al., 2016).

We hypothesize that chronic sleep disturbance, which is a risk factor for AD, would negatively affect the hippocampal neural plasticity important for pattern separation (Bekinschtein et al., 2014; Kent et al., 2015a; Kent et al., 2015b; Kent & Mistlberger, 2017). Indeed, a few studies in humans have shown that performance on tests designed to assess pattern separation are sensitive to periods of wakefulness and sleep deprivation in young adults (Cellini, 2023; Doxey et al., 2018; Hanert et al., 2017; Saletin et al., 2016). Thus, cognitive assessments designed to assess pattern separation have been shown to be sensitive to both early stages of AD (Stark et al., 2019) and sleep (Cellini, 2023; Doxey et al., 2018; Hanert et al., 2017; Saletin et al., 2016).

Here, we conducted an exploratory study examining the relationship between sleep and performance on two cognitive tests that tax pattern separation processes, the Mnemonic Similarity Task (MST) (Stark et al., 2019) and the Delayed Matching to Sample (DMS) test (Cambridge Cognition, 2023a), which is part of the Cambridge Neuropsychological Test Automated Battery (CANTAB). We compared performance on these tests of pattern separation to the Psychomotor Vigilance Task (PVT), which has been previously shown to be sensitive to sleep loss and circadian misalignment (Dinges & Powell, 1985; Lim & Dinges, 2008) as well as performance on the other CANTAB tests that are part of the Prodromal Alzheimer’s and MCI battery. We hypothesized that objective shorter sleep duration and poorer sleep quality would be associated with longer reaction times on the PVT, and more memory errors on the MST and CANTAB DMS.

## 2. Materials and Methods

### 2.1 Participants

The study was approved by the Research Ethics Board (REB) at Simon Fraser University (Protocol #30000539) in compliance with the Canadian Tri Council Policy Statement: Ethical Conduct of Research Involving Humans. Participants were adults 50+ years of age, recruited via posters, word-of-mouth, social media platforms, and the online platform REACHBC (https://reachbc.ca/). Prior to any testing, participants completed an online consent form and a questionnaire, which collected demographic and health information, such as information regarding sleep disorders. Participants needed to be able to understand and follow written and verbal instructions to be eligible to participate. There were no other exclusion criteria. Upon completion of the study, the participants were compensated with payment.

### 2.2 Procedure

Participants were scheduled for two in-person visits. All visits took place between 10:00 and 15:00. During the first visit, participants were provided with instructions and wrist actigraphy. On the 7th day, participants returned to the lab to complete cognitive testing. Figure 1 provides an illustration of the procedure. Participants completed the tasks in the same order: PVT, MST, CANTAB, and then MoCA. The testing took ∼75 min to complete.

**Figure 1.**
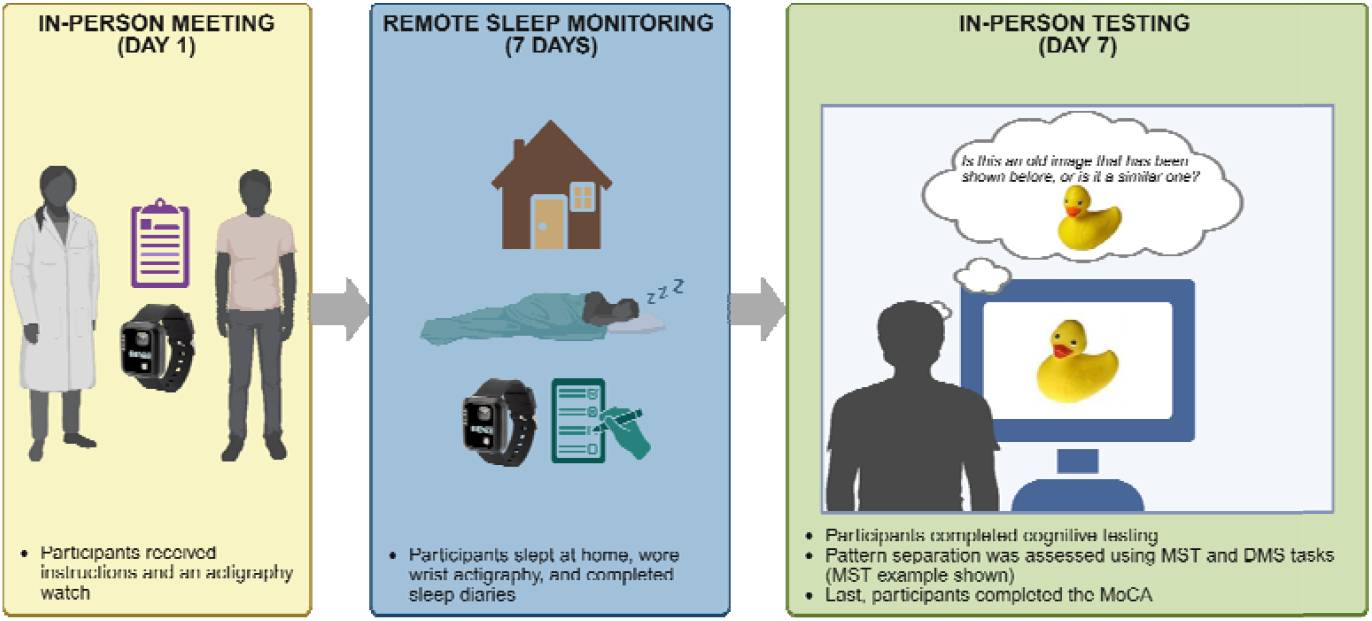
Illustration of the procedure, including two example images used in the MST (adapted from Stark et al., 2019). Created with BioRender.com. Abbreviations: MST, Mnemonic Similarity Task; DMS, Delayed Matching to Sample.

### 2.3 Sleep Assessments

Sleep monitoring was performed by the participants in their home environment. All participants wore wrist actigraphy (ActTrust2 model AT20101, Condor Instruments, Sao Paulo, Brazil) on their non-dominant hand and completed an online sleep diary, hosted via Qualtrics (https://www.qualtrics.com/), for 7 consecutive days. Participants were asked to press an event button on the wrist actigraphy when going to bed and when waking up in the morning. The data were sampled in 1-min epochs. We asked the participants to follow their habitual sleep schedule and did not enforce a schedule.

### 2.4 Psychomotor Vigilance Task (PVT)

The PVT is designed to measure sustained attention and reaction time (Dinges & Powell, 1985; Wilkinson & Houghton, 1982). We used the 5-min PVT version developed by NASA (Version 1.2.0., Washington, D.C., USA) (Arsintescu et al., 2017, 2019). The task was presented on an Apple iPad (OS 15.4.1, model A2602). The participants were instructed to attend to a small rectangular area on the dark iPad screen where a millisecond counter appeared, rapidly increasing from zero. The millisecond counter appeared at random inter-stimulus intervals (ISI), ranging from 2 to 10 s (Arsintescu et al., 2017). The participants were asked to tap the thumb of their dominant hand on the screen as quickly as possible when the number was shown. A practice session was completed before the experimental trials.

### 2.5 Mnemonic Similarity Task (MST)

The MST (Version 0.96) was used to assess pattern separation (Stark et al., 2019). We used the 3-choice version and the pace of the task was computerized (i.e., not self-paced). The task was run using Set 1 stimuli and included two phases. Phase 1 had 128 trials and participants were sequentially presented with images of everyday objects and asked to indicate via keyboard responses whether the object was an indoor or outdoor object (Stark et al., 2023, 2019). The participants were not informed about the upcoming memory test while they performed phase 1 of the task. During phase 2, which had 192 trials, the participants were sequentially presented with images of everyday objects, and the participants were asked to respond “old,” “similar,” or “new” to indicate whether the objects were a) identical to an image shown in phase 1 (i.e old/target) b) similar to an image presented in phase 1 (i.e., similar/lure), or c) a new image not presented in phase 1 (i.e., new/foil). Of the 192 trials, one-third (64) were targets, one-third (64) were lures, and one-third (64) were foils. The lures varied by 5 levels of similarity ranging from L1 (highly similar) to L5 (least similar). All images in both phases were presented with a duration of 2000 ms with an inter-stimulus interval of 500 ms.

### 2.6 Cambridge Neuropsychological Test Automated Battery (CANTAB)

The Prodromal Alzheimer’s and MCI CANTAB battery included 4 cognitive tests: Delayed Matching to Sample (DMS), Paired Associates Learning (PAL), Reaction Time (RTI), and Spatial Working Memory (SWM). All tests were presented on an Apple iPad (OS 15.4.1).

The Delayed Matching to Sample (DMS) test measured visual matching ability and short-term visual recognition memory for non-verbalizable patterns. During the task, participants were shown a complex pattern (i.e., sample pattern) for 4.5 s, and four choice patterns, composed of different configurations and colors. One of the choice patterns was identical to the sample pattern and participants were asked to select one that matched. In some trials, the choice patterns and the sample pattern appeared simultaneously, whereas in other trials, there was a delay of 0, 4, or 12 s before the four choices were presented (Cambridge Cognition, 2023a).

The Paired Associates Learning (PAL) task assessed new learning and visual episodic memory (Cambridge Cognition, 2023b). During the task, boxes were shown on the screen and were opened in a randomized order for 2 s. One or more of the boxes contained a pattern. The patterns were then presented in the center of the screen, one at a time. The participant was asked to select the box (i.e., location) in which the pattern was originally displayed.

The Reaction Time (RTI) task assessed divided attention and psychomotor speed (Cambridge Cognition, 2023c). During the task, the participant was asked to select and hold down a button at the bottom of the screen to make a yellow flash appear in one of five circles presented at the top of the screen. Once the flash appeared, the participant was instructed to react as quickly as possible by releasing the button at the bottom of the screen and touching the circle in which the yellow flash appeared.

The Spatial Working Memory (SWM) task required manipulation and retention of visuospatial information (Cambridge Cognition, 2023d). During this task, multiple boxes were presented on the screen, with one containing a hidden target. The participant was asked to select the empty boxes until only one box remained that contained the hidden target.

### 2.7 The Montreal Cognitive Assessment (MoCA)

The participants completed the MoCA Duo (MoCA Montreal) administered on an Apple iPad (OS 15.4.1, model A2602 with a 260.6 × 174.1 mm screen size) (Nasreddine et al., 2005). MoCA Duo is an automated version of the MoCA (MoCA Cognition, 2024). The MoCA measures 6 cognitive domains including executive function, memory, attention, visuospatial skills, language, and orientation (Nasreddine et al., 2005). The test is a 30-point scale and a total score of less than 26 can indicate risk of MCI or dementia. An extra point is added for participants who have fewer than 12 years of formal education. The administration time of the test was approximately 10 min. MoCA was used to evaluate global cognition and not used to screen the participants.

### 2.8 Data analysis

#### 2.8.1 Wrist actigraphy

Wrist actigraphy data were extracted and analyzed using ActStudio software (Version 2.2.0, Condor Instruments, Sao Paulo, Brazil). Only nocturnal sleep was assessed. Average total sleep time (TST) across the week was used as a measure of sleep quantity and average sleep efficiency across the week was used as a measure of sleep quality. These measures are commonly used sleep metrics in research in older adults (Ghazi et al., 2025) and tend to decline in aging (Li et al., 2018) and in those at risk for cognitive decline and dementia (Kent et al., 2021). A minimum of 5 nights of complete wrist actigraphy data were required for each participant to be included in the analyses. Sleep diaries were used to confirm evening bedtimes and morning wake-up times. We also used Clocklab (Version 6.1.11, Actimetrics, Wilmette, USA) to run a cosinor analysis to estimate acrophase, amplitude, interdaily stability (IS), and intradaily variability (IV) of the daily activity rhythms.

#### 2.8.2 PVT

We calculated the slowest 10% reaction times for all PVT trials, which has previously been shown to be sensitive to sleep loss (Basner & Dinges, 2011; Loh et al., 2004).

#### 2.8.3 MST

For the MST analyses, the Memory Recognition scores (REC) were calculated as [p(correct old response to repeated images) - p(incorrect old response to foils)] (Stark et al., 2019). The Lure Discrimination Index (LDI) was calculated as [p(correct similar response to lures) - p(incorrect similar response to foils)] (Stark et al., 2013). We also assessed performance for the L1 and L2 specific lure bins, which have the highest levels of similarity between the lures and the repeated images, and thus are most taxing on pattern separation. For L1 and L2, we assessed *Accuracy*, as the probability of correctly identifying the lure as similar, and *False Memory Error Rate*, as the probability of falsely identifying the lure as a repeated image.

#### 2.8.4 CANTAB

CANTAB data were analyzed using the CANTAB software (Cambridge Cognition, 2021). The key performance measures included in our analyses were:

##### DMS

We calculated DMS Pattern Errors (All Delays), as the number of times the participant initially chose the distractor stimulus with different physical attributes (i.e., an incorrect pattern), but matching color elements, calculated across all trials with a delay component.

##### PAL

The number of times a participant selected the correct box on their initial attempt while recalling the pattern locations (PAL First Attempt), and the number of times a participant selected the incorrect box with an adjustment to allow for a comparison of error rates across all participants (PAL Adjusted Errors).

##### RTI

The average duration elapsed from the moment a participant let go of the response button until they selected the target stimulus (RTI Mean Five-Choice Movement Time).

##### SWM

The number of times a participant revisited a box where a token was previously found (SWM Between Errors) and the number of times the participant started a new search pattern from the same initial box in trials with 6-8 boxes (SWM Strategy).

#### 2.8.5 Montreal Cognitive Assessment

The MoCA total score calculated out of 30 points.

#### 2.8.6 Statistical analyses

The relationships between sleep measures and cognitive performance were first assessed using an exploratory approach including Pearson’s correlation analyses, using JMP Student Edition 18.2.1 (JMP Statistical Discovery LLC, North Carolina, USA). Correlations between data that violated the assumption of normality were conducted using Spearman’s rank correlation (ρ). Significant correlations between sleep and cognitive performance were further examined using multiple linear regression analyses, to account for age and sex as potential confounding variables. We also ran an additional model controlling for the effect of education on performance. Due to the sample size, we did not include other covariates (e.g., sleep disorders) in the analyses. The residuals of the models were approximately normally distributed, centered around zero with constant variance and were not affected by outliers. Observations were independent shown by the Durbin Watson’s test (*p* > 0.05). We did not include ethnicity in our model as our sample size did not allow for equal distribution of different ethnic groups. The cut-off used for statistical significance was *p* < 0.05 (two-tailed) for all measures. As the correlations were used as an exploratory analysis, we applied the Holm-Bonferroni procedure to control for multiple comparisons only in the regression analyses.

## 3. Results

### 3.1. Sample characteristics

Of the 40 participants recruited, 6 participants were excluded from all analyses, due to unreliable or missing sleep data (n = 1), not understanding task instructions (n = 2), poor performance or malfunction on both MST and CANTAB (n = 3). The remaining 34 participants (mean age: 65.56 ± 9.57) were included in the analyses. Table 1 shows the participant demographic information and Table 2 shows the average sleep and circadian rhythm parameters.

**Table 1.**
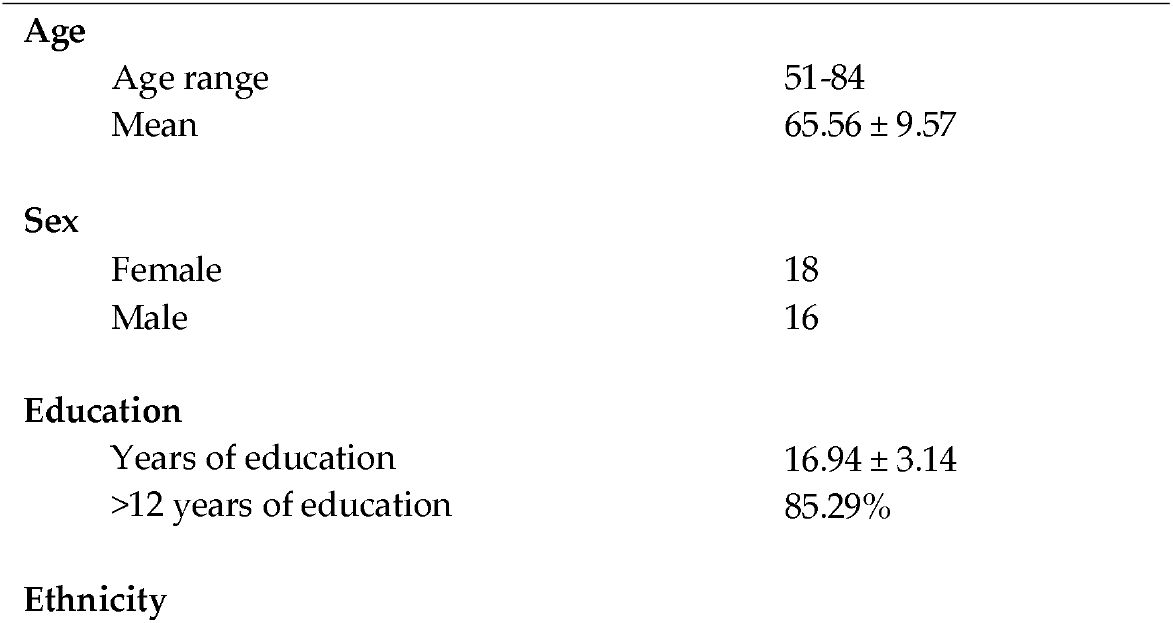
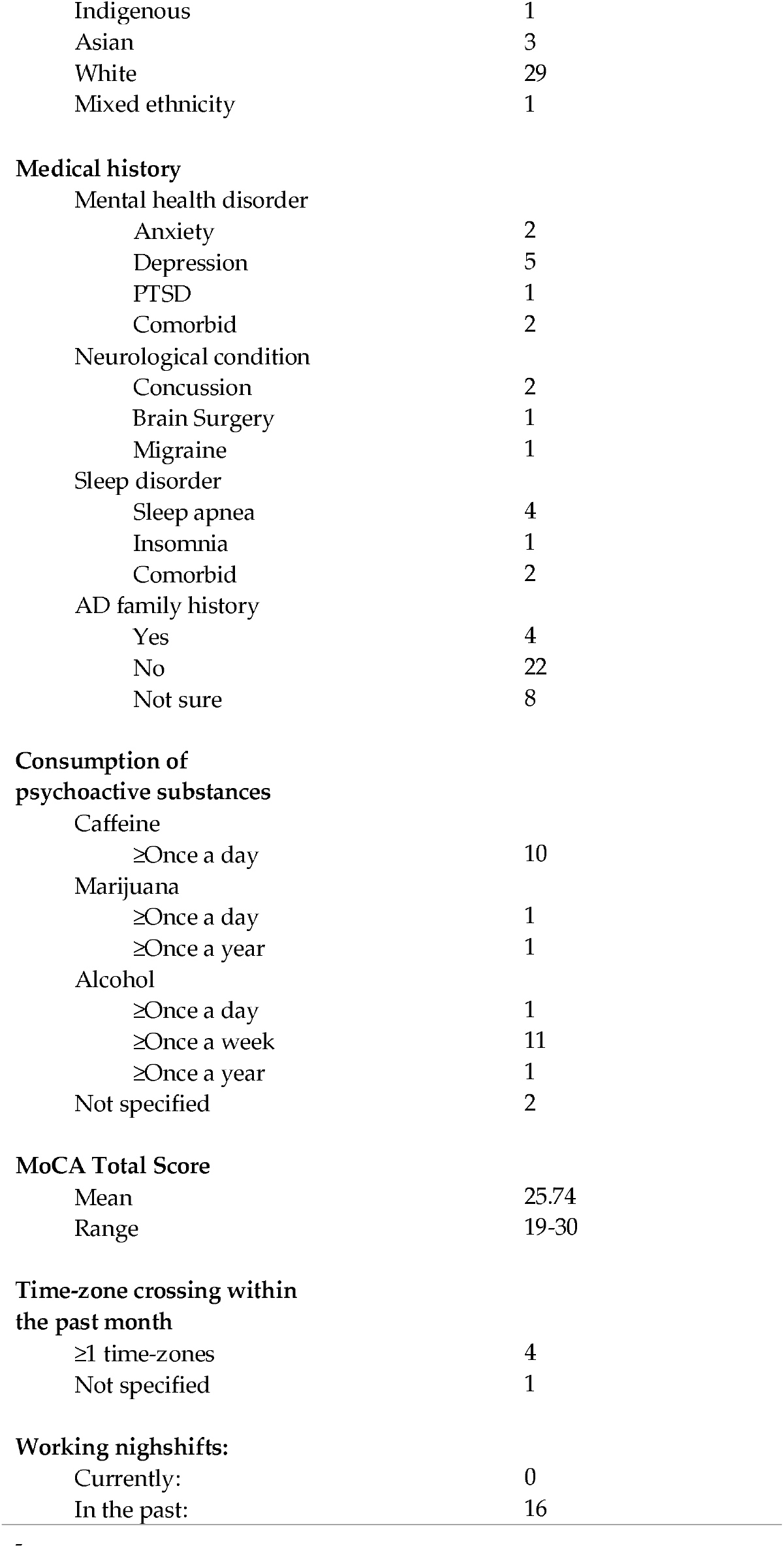
Participant characteristics.

|  |  |  |
| --- | --- | --- |
| **Age** |  |  |
|  | Age range | 51-84 |
| | Mean | $65.56 \pm 9.57$ |
| **Sex** |  |  |
|  | Female | 18 |
|  | Male | 16 |
| **Education** |  |  |
| | Years of education | $16.94 \pm 3.14$ |
|  | >12 years of education | 85.29% |
| **Ethnicity** |  |  |

|  |  |
| --- | --- |
| Indigenous | 1 |
| Asian | 3 |
| White | 29 |
| Mixed ethnicity | 1 |
| **Medical history** |  |
| Mental health disorder |  |
| Anxiety | 2 |
| Depression | 5 |
| PTSD | 1 |
| Comorbid | 2 |
| Neurological condition |  |
| Concussion | 2 |
| Brain Surgery | 1 |
| Migraine | 1 |
| Sleep disorder |  |
| Sleep apnea | 4 |
| Insomnia | 1 |
| Comorbid | 2 |
| AD family history |  |
| Yes | 4 |
| No | 22 |
| Not sure | 8 |
| **Consumption of psychoactive substances** |  |
| Caffeine |  |
| ≥Once a day | 10 |
| Marijuana |  |
| ≥Once a day | 1 |
| ≥Once a year | 1 |
| Alcohol |  |
| ≥Once a day | 1 |
| ≥Once a week | 11 |
| ≥Once a year | 1 |
| Not specified | 2 |
| **MoCA Total Score** |  |
| Mean | 25.74 |
| Range | 19-30 |
| **Time-zone crossing within the past month** |  |
| ≥1 time-zones | 4 |
| Not specified | 1 |
| **Working nighshifts:** |  |
| Currently: | 0 |
| In the past: | 16 |

**Table 2.** Summary of sleep and circadian rhythm parameters.

|  | Bedtime | Wake-up<br>time | TST (h) | Sleep efficiency<br>(%) | SOL (h) | WASO (h) | Amplitude | Acrophase<br>(h) | IV | IS |
| --- | --- | --- | --- | --- | --- | --- | --- | --- | --- | --- |
|  | Mean ± SD | Mean ± SD | Mean ± SD | Mean ± SD | Mean ± SD | Mean ± SD | Mean ± SD | Mean ± SD | Mean ± SD | Mean ± SD |
| Participants | 23.08 ± 1.38 | 7.01 ± 1.22 | 7.18 ± 0.98 | 90.55 ± 4.21 | 0.06 ± 0.02 | 0.65 ± 0.34 | 1.68 ± 0.66 | 14.30 ± 1.97 | 1.14 ± 0.31 | 0.42 ± 0.14 |

Abbreviations: TST, Total Sleep Time; SOL, Sleep Onset Latency; WASO, Wake After Sleep Onset; IV, Intradaily Variability; IS, Interdaily Stability. Bedtime, Wake-up time, TST, SOL, WASO, and acrophase are displayed in decimal time.

For the task-specific analyses, an additional 4 participants were excluded from PVT analyses due to reporting major distractions during task performance (n = 1) or failing to keep their thumb still in between trials, leading to unintentional screen taps (n = 3). Nine participants were excluded from MST analyses due to mixing up button presses (n = 1), not understanding the task indicated by using the similar response less than 10 times (n = 5) (Callow et al., 2023), not complying with task instructions (n = 1), and a REC score below the cut-off value of 0.50 (n = 2). No participants were excluded from CANTAB analyses. Two participants were excluded from circadian analyses as they did not have 5 or more consecutive days of continuous wrist actigraphy-wearing.

### 3.2 Performance on PVT is correlated with sleep

There were significant correlations between average TST and mean slowest 10% RT (*ρ*[28] = -0.38, *p* = 0.04). There were no significant correlations between average sleep efficiency and PVT performance (*p* > 0.05); See Table 3. There were also no significant correlations between PVT and circadian parameters (*p* > 0.05; See Table 4).

**Table 3.**
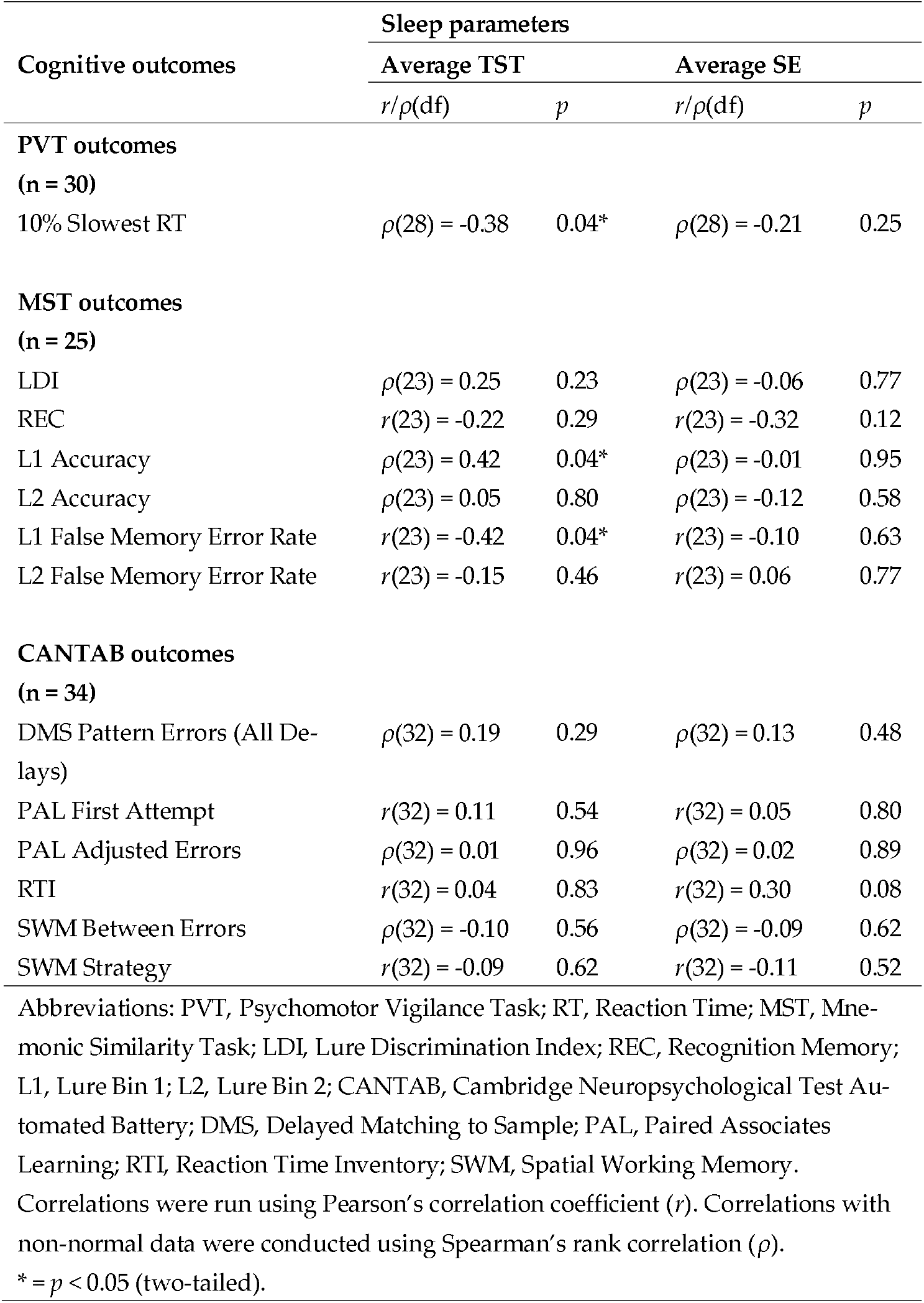
Correlations between total sleep time (TST), sleep efficiency (SE), and performance on cognitive tests.

**Table 4.**
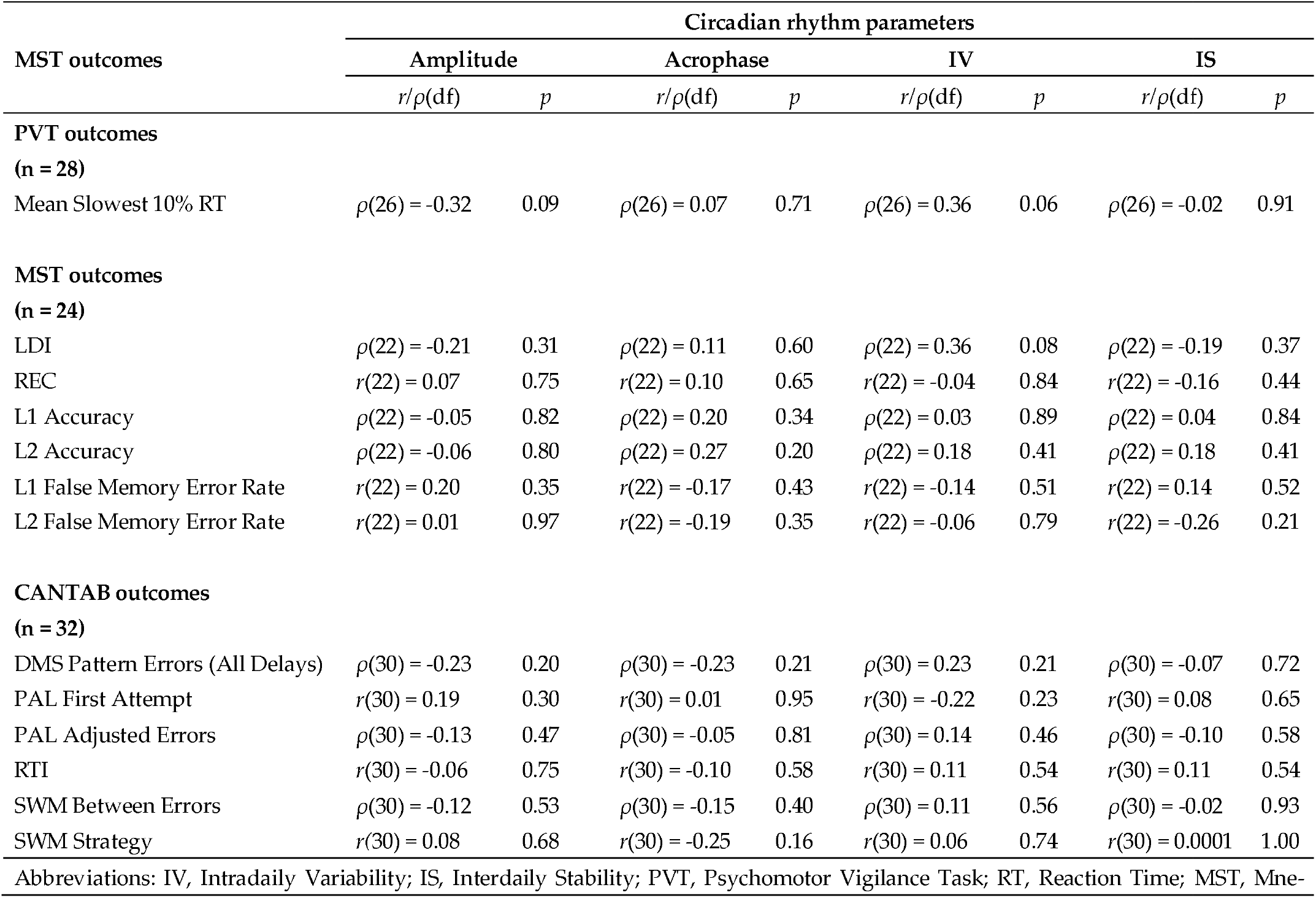
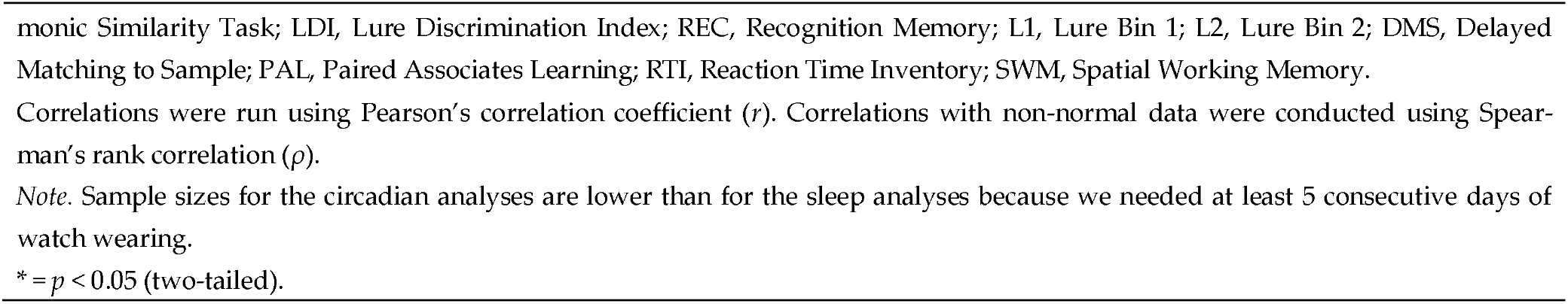
Correlations between performance on cognitive tests and circadian rhythm parameters.

| MST outcomes | Circadian rhythm parameters |  |  |  |  |  |  |  |
| --- | --- | --- | --- | --- | --- | --- | --- | --- |
|  | Amplitude |  | Acrophase |  | IV |  | IS |  |
| | $r/\rho(df)$ | $p$ | $r/\rho(df)$ | $p$ | $r/\rho(df)$ | $p$ | $r/\rho(df)$ | $p$ |
| **PVT outcomes** |  |  |  |  |  |  |  |  |
| **(n = 28)** |  |  |  |  |  |  |  |  |
| Mean Slowest 10% RT | $\rho(26) = -0.32$ | 0.09 | $\rho(26) = 0.07$ | 0.71 | $\rho(26) = 0.36$ | 0.06 | $\rho(26) = -0.02$ | 0.91 |
| **MST outcomes** |  |  |  |  |  |  |  |  |
| **(n = 24)** |  |  |  |  |  |  |  |  |
| LDI | $\rho(22) = -0.21$ | 0.31 | $\rho(22) = 0.11$ | 0.60 | $\rho(22) = 0.36$ | 0.08 | $\rho(22) = -0.19$ | 0.37 |
| REC | $r(22) = 0.07$ | 0.75 | $r(22) = 0.10$ | 0.65 | $r(22) = -0.04$ | 0.84 | $r(22) = -0.16$ | 0.44 |
| L1 Accuracy | $\rho(22) = -0.05$ | 0.82 | $\rho(22) = 0.20$ | 0.34 | $\rho(22) = 0.03$ | 0.89 | $\rho(22) = 0.04$ | 0.84 |
| L2 Accuracy | $\rho(22) = -0.06$ | 0.80 | $\rho(22) = 0.27$ | 0.20 | $\rho(22) = 0.18$ | 0.41 | $\rho(22) = 0.18$ | 0.41 |
| L1 False Memory Error Rate | $r(22) = 0.20$ | 0.35 | $r(22) = -0.17$ | 0.43 | $r(22) = -0.14$ | 0.51 | $r(22) = 0.14$ | 0.52 |
| L2 False Memory Error Rate | $r(22) = 0.01$ | 0.97 | $r(22) = -0.19$ | 0.35 | $r(22) = -0.06$ | 0.79 | $r(22) = -0.26$ | 0.21 |
| **CANTAB outcomes** |  |  |  |  |  |  |  |  |
| **(n = 32)** |  |  |  |  |  |  |  |  |
| DMS Pattern Errors (All Delays) | $\rho(30) = -0.23$ | 0.20 | $\rho(30) = -0.23$ | 0.21 | $\rho(30) = 0.23$ | 0.21 | $\rho(30) = -0.07$ | 0.72 |
| PAL First Attempt | $r(30) = 0.19$ | 0.30 | $r(30) = 0.01$ | 0.95 | $r(30) = -0.22$ | 0.23 | $r(30) = 0.08$ | 0.65 |
| PAL Adjusted Errors | $\rho(30) = -0.13$ | 0.47 | $\rho(30) = -0.05$ | 0.81 | $\rho(30) = 0.14$ | 0.46 | $\rho(30) = -0.10$ | 0.58 |
| RTI | $r(30) = -0.06$ | 0.75 | $r(30) = -0.10$ | 0.58 | $r(30) = 0.11$ | 0.54 | $r(30) = 0.11$ | 0.54 |
| SWM Between Errors | $\rho(30) = -0.12$ | 0.53 | $\rho(30) = -0.15$ | 0.40 | $\rho(30) = 0.11$ | 0.56 | $\rho(30) = -0.02$ | 0.93 |
| SWM Strategy | $r(30) = 0.08$ | 0.68 | $r(30) = -0.25$ | 0.16 | $r(30) = 0.06$ | 0.74 | $r(30) = 0.0001$ | 1.00 |
| Abbreviations: IV, Intradaily Variability; IS, Interdaily Stability; PVT, Psychomotor Vigilance Task; RT, Reaction Time; MST, Mne- |  |  |  |  |  |  |  |  |

monic Similarity Task; LDI, Lure Discrimination Index; REC, Recognition Memory; L1, Lure Bin 1; L2, Lure Bin 2; DMS, Delayed Matching to Sample; PAL, Paired Associates Learning; RTI, Reaction Time Inventory; SWM, Spatial Working Memory.
Correlations were run using Pearson's correlation coefficient ( $r$ ). Correlations with non-normal data were conducted using Spearman's rank correlation ( $\rho$ ).
*Note.* Sample sizes for the circadian analyses are lower than for the sleep analyses because we needed at least 5 consecutive days of watch wearing.
\* = $p < 0.05$ (two-tailed).

### 3.3 Performance on a cognitive test of pattern separation is correlated with sleep

There were statistically significant correlations between sleep and MST performance; specifically, TST and L1 Accuracy (*ρ*[23] = 0.42, *p* = 0.04) (Figure 2c), and between TST and L1 False Memory Error Rate (r[23] = -0.42, *p* = 0.04) (Figure 2e). There were no statistically significant correlations between sleep and performance on the CANTAB battery (*p* > 0.05). See Table 3 for the remaining results. There were also no significant relationships between MST or CANTAB tests and circadian parameters (*p* > 0.05). Remaining statistical results are displayed in Table 4.

**Figure 2.**
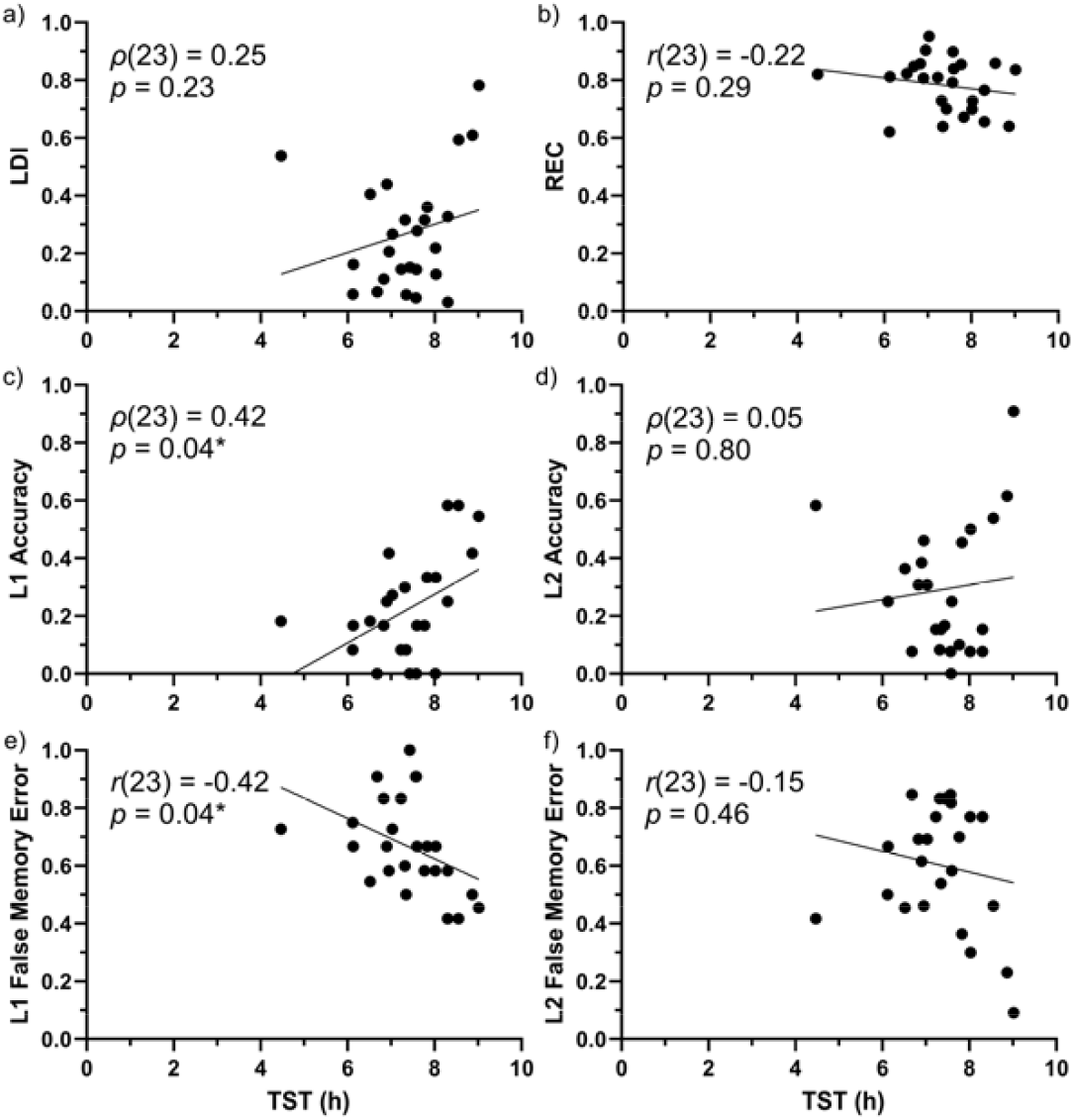
Scatterplots showing the relationship between TST and MST performance: LDI (a), REC (b), L1 Accuracy (c), L2 Accuracy (d), L1 False Memory Error Rate (e), and L2 False Memory Error Rate (f). Each black dot represents an individual participant. Pearson’s (r) or Spearman’s (ρ) correlation, ^*^ = p < 0.05. Abbreviations: TST, total sleep time; LDI, Lure Discrimination Index; REC, Recognition Memory; L1, Lure Bin 1; L2, Lure Bin 2.

### 3.4 MoCA performance was not correlated with sleep or circadian parameters but was correlated with cognitive performance measures

There were no relationships between the MoCA total score and TST or sleep efficiency, or circadian parameters (*p* > 0.05). The PVT was also not significantly correlated with the MoCA (*p* > 0.05). For the MST, the MoCA total score was significantly associated with L1 Accuracy (*ρ*[23] = 0.45, *p* = 0.02) (Figure 3c) and L1 False Memory Error Rate (r[23] = -0.42, *p* = 0.04) (Figure 3e). For CANTAB, the MoCA total score was significantly correlated with PAL First Attempt (r[32] = 0.51, *p* = 0.002), PAL Adjusted Errors (r[32] = -0.42, *p* = 0.01), and the SWM measure SWM Between Errors (*ρ*[32] = -0.53, *p* = 0.001). See Table 5 for the remaining results.

**Table 5.**
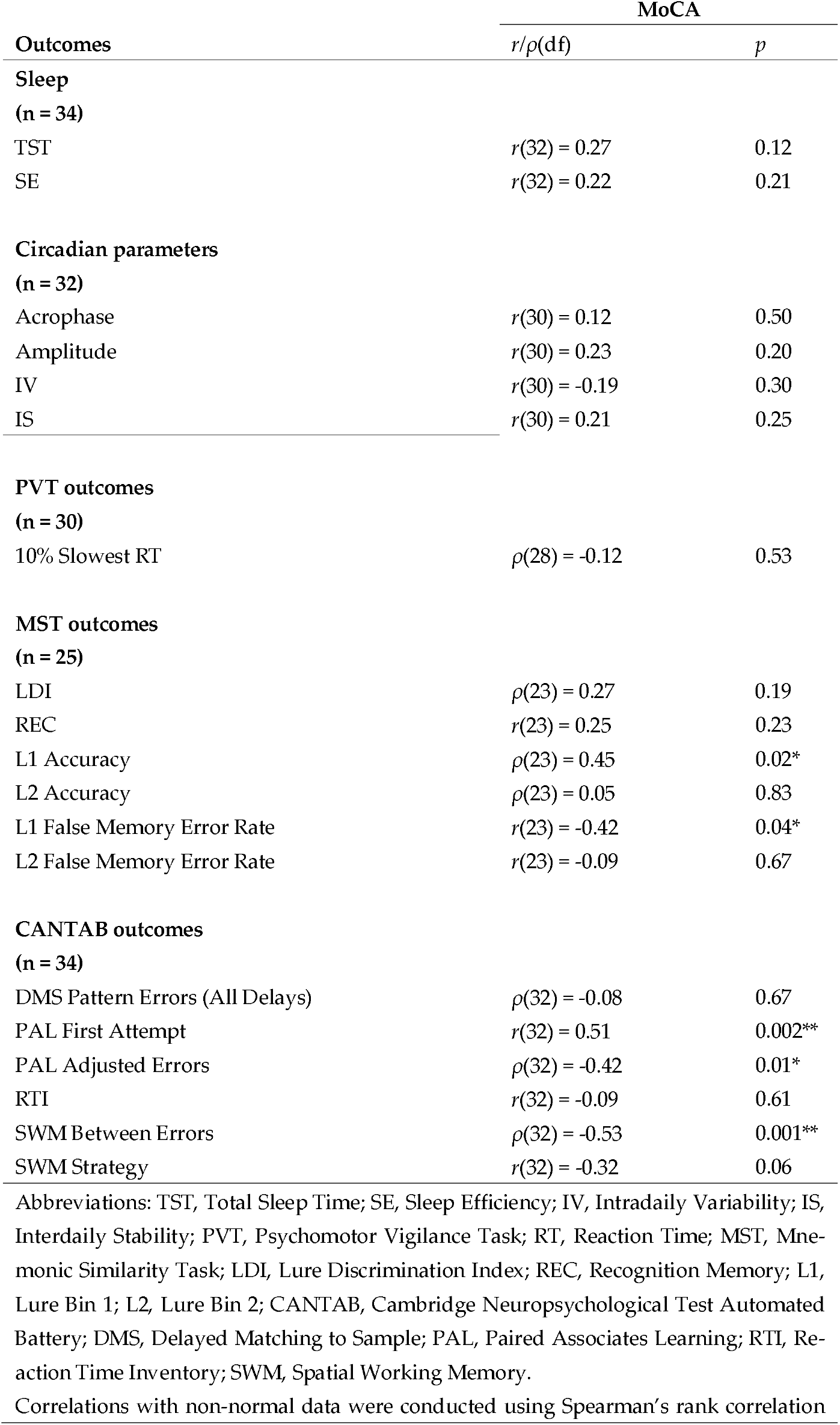
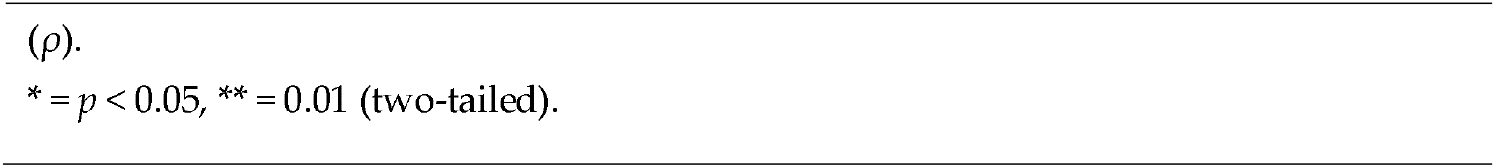
Correlations between sleep and circadian rhythm outputs, performance on cognitive tests, and the MoCA.

| cognitive tests, and the MoCA |  |  |
| --- | --- | --- |
| Outcomes | MoCA |  |
| | $r/\rho(df)$ | $p$ |
| **Sleep** |  |  |
| **(n = 34)** |  |  |
| TST | $r(32) = 0.27$ | 0.12 |
| SE | $r(32) = 0.22$ | 0.21 |
| **Circadian parameters** |  |  |
| **(n = 32)** |  |  |
| Acrophase | $r(30) = 0.12$ | 0.50 |
| Amplitude | $r(30) = 0.23$ | 0.20 |
| IV | $r(30) = -0.19$ | 0.30 |
| IS | $r(30) = 0.21$ | 0.25 |
| **PVT outcomes** |  |  |
| **(n = 30)** |  |  |
| 10% Slowest RT | $\rho(28) = -0.12$ | 0.53 |
| **MST outcomes** |  |  |
| **(n = 25)** |  |  |
| LDI | $\rho(23) = 0.27$ | 0.19 |
| REC | $r(23) = 0.25$ | 0.23 |
| L1 Accuracy | $\rho(23) = 0.45$ | 0.02* |
| L2 Accuracy | $\rho(23) = 0.05$ | 0.83 |
| L1 False Memory Error Rate | $r(23) = -0.42$ | 0.04* |
| L2 False Memory Error Rate | $r(23) = -0.09$ | 0.67 |
| **CANTAB outcomes** |  |  |
| **(n = 34)** |  |  |
| DMS Pattern Errors (All Delays) | $\rho(32) = -0.08$ | 0.67 |
| PAL First Attempt | $r(32) = 0.51$ | 0.002** |
| PAL Adjusted Errors | $\rho(32) = -0.42$ | 0.01* |
| RTI | $r(32) = -0.09$ | 0.61 |
| SWM Between Errors | $\rho(32) = -0.53$ | 0.001** |
| SWM Strategy | $r(32) = -0.32$ | 0.06 |

Abbreviations: TST, Total Sleep Time; SE, Sleep Efficiency; IV, Intradaily Variability; IS, Interdaily Stability; PVT, Psychomotor Vigilance Task; RT, Reaction Time; MST, Mnemonic Similarity Task; LDI, Lure Discrimination Index; REC, Recognition Memory; L1, Lure Bin 1; L2, Lure Bin 2; CANTAB, Cambridge Neuropsychological Test Automated Battery; DMS, Delayed Matching to Sample; PAL, Paired Associates Learning; RTI, Reaction Time Inventory; SWM, Spatial Working Memory.
Correlations with non-normal data were conducted using Spearman's rank correlation
\* = $p < 0.05$ , \*\* = 0.01 (two-tailed).

**Figure 3.**
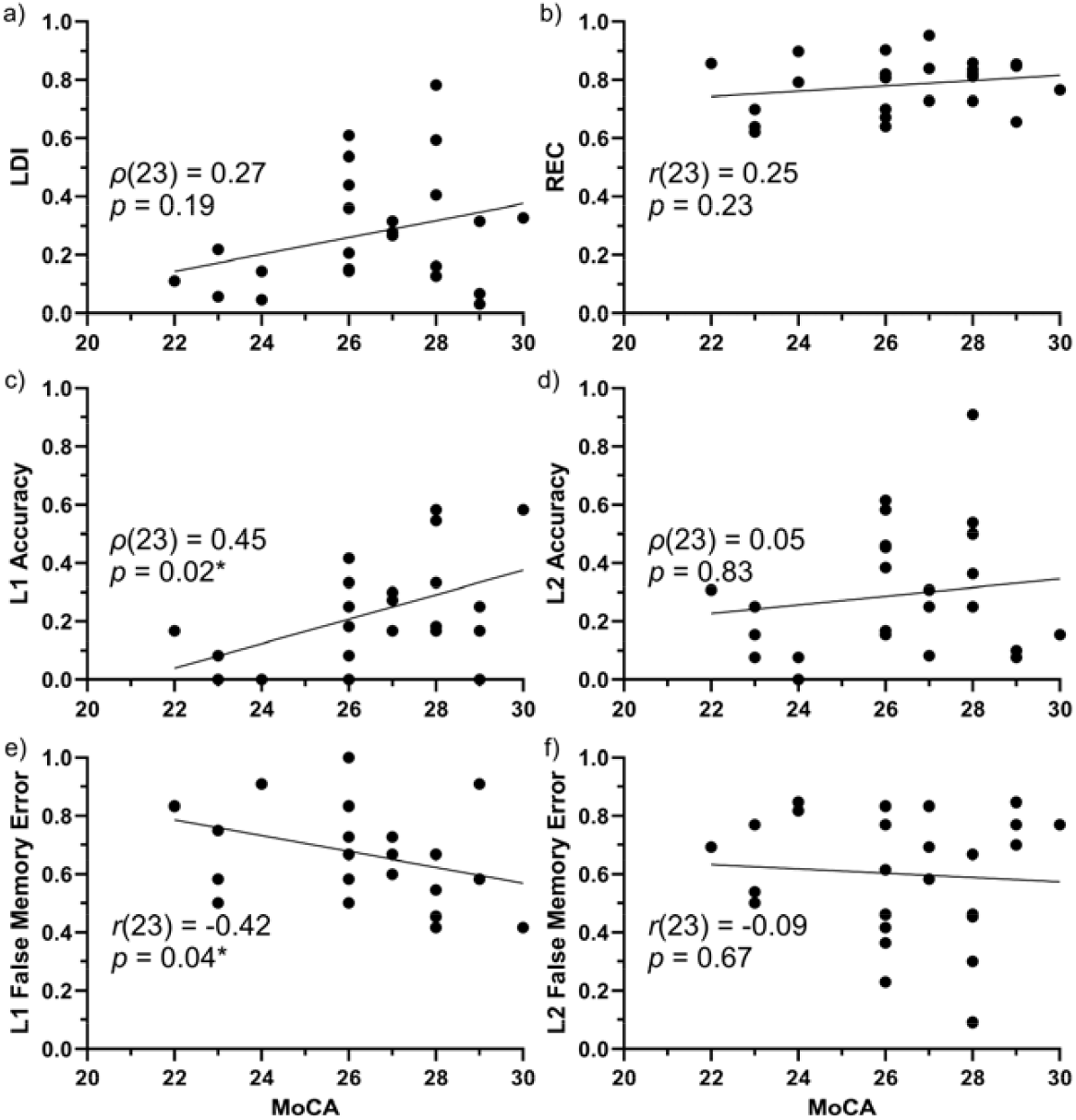
Relationship between performance on MoCA and MST. Scatterplots showing the relationship between MoCA total score and a) LDI, b) REC, c) L1 Accuracy, d) L2 Accuracy, e) L1 False Memory Error Rate, and f) L2 False Memory Error Rate. Each black dot represents an individual participant. Pearson’s (r) or Spearman’s (ρ) correlation, ^*^ = *p* < 0.05. Abbreviations: MoCA, Montreal Cognitive Assessment; LDI, Lure Discrimination Index; REC, Recognition Memory; L1, Lure Bin 1; L2, Lure Bin 2.

### 3.5 Performance on a cognitive test of pattern separation is predicted by total sleep time

When controlling for age and sex, the multiple linear regressions showed a significant overall model for L1 Accuracy (F_3,21_ = 10.65, *p* = 0.0002) and L1 False Memory Error Rate (F_3,21_ = 4.60, *p* = 0.01). Age was a significant covariate for L1 Accuracy (t_1,21_ = -4.52, *p* = 0.0002) and for L1 False Memory Error Rate (t_1,21_ = 2.75, *p* = 0.01). TST predicted L1 Accuracy (t_1,21_ = 3.34, *p* = 0.003) and L1 False Memory Error Rate (t_1,21_ = -2.95, *p* = 0.008). See Table 6 for additional statistical results. When applying the Holm-Bonferroni correction, age still predicted L1 Accuracy and TST remained a significant predictor in both models when using a threshold of α_adj_ = 0.0167. A sensitivity analysis including participants who had data from all measures (n=22) supported these findings, and age also significantly predicted L1 False Memory Error Rate in the sensitivity analysis when Holm-Bonferroni correction was applied (see Supplementary Table S1).

**Table 6.**
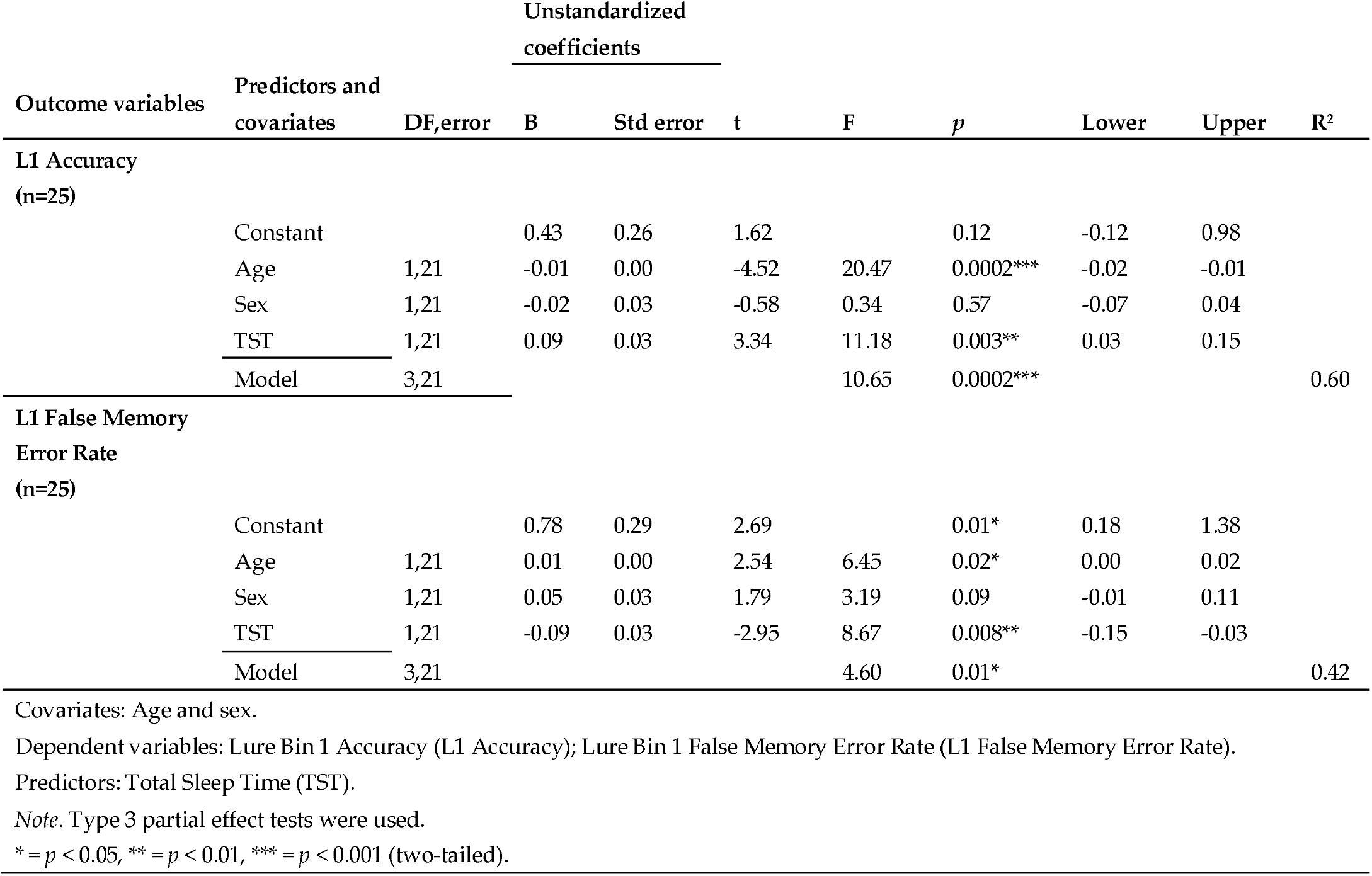
Multiple linear regression on the effect of total sleep time (TST) on MST performance.

| Outcome variables | Predictors and<br>covariates | DF,error | Unstandardized<br>coefficients |  | t | F | p | Lower | Upper | R<sup>2</sup> |
| --- | --- | --- | --- | --- | --- | --- | --- | --- | --- | --- |
|  |  |  | B | Std error |  |  |  |  |  |  |
| L1 Accuracy<br>(n=25) |  |  |  |  |  |  |  |  |  |  |
|  | Constant |  | 0.43 | 0.26 | 1.62 |  | 0.12 | -0.12 | 0.98 |  |
|  | Age | 1,21 | -0.01 | 0.00 | -4.52 | 20.47 | 0.0002*** | -0.02 | -0.01 |  |
|  | Sex | 1,21 | -0.02 | 0.03 | -0.58 | 0.34 | 0.57 | -0.07 | 0.04 |  |
|  | TST | 1,21 | 0.09 | 0.03 | 3.34 | 11.18 | 0.003** | 0.03 | 0.15 |  |
|  | Model | 3,21 |  |  |  | 10.65 | 0.0002*** |  |  | 0.60 |
| L1 False Memory<br>Error Rate<br>(n=25) |  |  |  |  |  |  |  |  |  |  |
|  | Constant |  | 0.78 | 0.29 | 2.69 |  | 0.01* | 0.18 | 1.38 |  |
|  | Age | 1,21 | 0.01 | 0.00 | 2.54 | 6.45 | 0.02* | 0.00 | 0.02 |  |
|  | Sex | 1,21 | 0.05 | 0.03 | 1.79 | 3.19 | 0.09 | -0.01 | 0.11 |  |
|  | TST | 1,21 | -0.09 | 0.03 | -2.95 | 8.67 | 0.008** | -0.15 | -0.03 |  |
|  | Model | 3,21 |  |  |  | 4.60 | 0.01* |  |  | 0.42 |

Covariates: Age and sex.
Dependent variables: Lure Bin 1 Accuracy (L1 Accuracy); Lure Bin 1 False Memory Error Rate (L1 False Memory Error Rate).
Predictors: Total Sleep Time (TST).
Note. Type 3 partial effect tests were used.
\* = $p < 0.05$ , \*\* = $p < 0.01$ , \*\*\* = $p < 0.001$ (two-tailed).

When education was added to the regression models, age (t_1,20_ = -4.40, p = 0.0003) and TST (t_1,20_ = 3.05, p = 0.006) remained significant predictors of L1 Accuracy when Holm-Bonferroni was applied. Age and TST did not significantly predict L1 False Memory Error Rate after controlling for multiple comparisons using a Holm-Bonferroni threshold of α_adj_ = 0.0125 (see Supplementary Table S2).

## 4. Discussion

It is well-established that sleep is important for optimal cognition and that sleep disturbances are often associated with psychiatric and neurological conditions that affect cognition. Recently, sleep has garnered attention as a potential therapeutic target for AD (Kent et al., 2021). Here, we conducted an exploratory study aiming to identify the cognitive tests that are most sensitive to sleep quantity and quality, focusing on those tests shown to be sensitive to the early cognitive changes associated with dementia. Clinical trials evaluating the effectiveness of sleep-promoting interventions in older adults at risk for or diagnosed with dementia require cognitive tests that are sensitive to sleep to be used as clinical trial outcome measures (Eyob et al., 2024).

We hypothesized that performance on tests designed to assess pattern separation (i.e., MST and CANTAB DMS) would be most sensitive to sleep quantity and quality (Kent & Mistlberger, 2017). The hallmark symptom of AD is impaired episodic memory. Importantly, failure in episodic memory does not always reflect forgetting, but instead the specific memory impairment exhibited by patients with MCI or AD may reflect false memory, because episodic memory is particularly vulnerable to interference (Budson et al., 2001; Hart et al., 1985). Pattern separation is a computational mechanism that plays an integral part in decreasing interference amongst similar memory representations (Kent et al., 2016). In the MST task, a failure in indicating lures as “similar” and instead judging lure images as “old” (false memory) may suggest ineffective pattern separation during encoding. Correctly judging lures as “similar” would indicate that effective pattern separation had taken place during encoding (Kirwan & Stark, 2007). In the DMS task, the participant has to differentiate between patterns that vary in similarity, thus, vary in the need for pattern separation. We hypothesize that the hippocampal neural plasticity required for effective pattern separation is sensitive to acute and chronic sleep loss, which makes these cognitive tests uniquely sensitive to sleep-associated cognition in older adults (Bekinschtein et al., 2014; Kent et al., 2015a; Kent et al., 2015b; Kent & Mistlberger, 2017).

Our results show that performance on MST is associated with sleep quantity in older adults. Specifically, we found a significant correlation between longer sleep duration in the participants and better discriminative performance on MST trials with the highest level of similarity between the lures and target images (i.e., L1). We also examined the probability of false memory on the MST lure trials with the highest similarity. We found that the participants with the longest sleep duration were less likely to have a false memory (i.e., falsely recognize an L1 Lure as a repeated image). Additionally, when controlling for age and sex, sleep duration predicted performance on MST lure trials with the highest similarity.

Previous research on sleep and pattern separation examined the effects of total sleep deprivation (Saletin et al., 2016), daytime wakefulness (Cellini, 2023; Doxey et al., 2018), and normal sleep (Cellini, 2023; Doxey et al., 2018) on LDI performance, and daytime wakefulness and normal sleep on lure bin performance (Hanert et al., 2017). For example, Saletin et al. (2016) showed that total sleep deprivation impairs LDI performance (Saletin et al., 2016), and Hanert et al. (2017) found that extended daytime wakefulness affects performance on the lure trials with the least similarity, whereas nighttime sleep improves performance on these trials (Hanert et al., 2017). To our knowledge, our study is the first to show that performance on images most similar to target stimuli are affected by varying sleep durations in older adults and the average performance of our cohort is well-aligned with previously reported MST performance of older adults, specifically (Stark et al., 2013, 2019). Comparing our participants’ mean age-related performance (mean age: 65.56) on MST LDI, our participants’ score (LDI = .27) closely aligns with that of participants’ LDI mean score performance of an approximately similar mean age in Stark et al. (2019) (see Figure 2E in Stark et al., 2019).

Performance on the CANTAB tests that are part of the Prodromal AD and MCI battery was not correlated with sleep. Most of the CANTAB tests, however, were correlated with MoCA total score, specifically PAL and SWM. This is consistent with previous studies showing the high sensitivity of PAL at detecting the earliest cognitive changes associated with dementia (Robbins & Sahakian, 2016). It also importantly highlights that not all tasks validated for detecting the earliest cognitive changes associated with MCI and dementia are equally as sensitive to sleep-associated cognitive performance.

Our results also show significant relationships between performance on MoCA and MST. Specifically, lower MoCA scores were associated with poorer performance on MST trials with the highest level of similarity between the lures and repeated images (i.e., L1). Our results complement but do not replicate previous work that showed that the MoCA score predicted LDI performance (Pishdadian et al., 2020). In our data, there does appear to be a non-significant association between the MoCA score and LDI, which might suggest that our study was unpowered to detect this relationship. Since L1 lures tax pattern separation the most, our analyses suggest that L1 lure discrimination and false memory errors may be more sensitive than overall LDI to dementia-related decline and sleep-related impairments. These findings support previous reports that MST is sensitive to AD-associated cognitive decline (Stark et al., 2013) and may be an effective outcome measure in clinical trials interested in measuring memory improvement associated with sleep interventions in patients with dementia (Bakker et al., 2012). Importantly, our results highlight that while both PAL and MST are correlated with MoCA scores, only MST (and not PAL) was also correlated with and predicted by sleep quantity.

Performance on the PVT was also associated with sleep quantity. While the PVT is sensitive to sleep, it was not correlated with MoCA scores or other cognitive measures indicative of dementia. The ideal clinical outcome measure for sleep targeting therapeutics for AD would be a cognitive measure sensitive to both sleep and AD-associated cognitive decline.

The current research had strengths. We used both wrist actigraphy and self-report measures to achieve an estimation of rest and activity. We also used cognitive tests that have been validated to assess cognitive domains known to decline in prodromal AD and MCI. To our knowledge, this is the first study examining the relationship between sleep quantity, sleep quality, and pattern separation in older adults using actigraphy watches. Previous studies on sleep and pattern separation performance recruited healthy younger adults (Cellini, 2023; Doxey et al., 2018; Hanert et al., 2017; Saletin et al., 2016), and one study implemented wrist actigraphy as sleep measure (Doxey et al., 2018).

The current research also had limitations. Our study did not include direct measures of sleep or measures that allowed for sleep staging (e.g., polysomnography or at-home EEG). Additionally, caffeine intake may have affected performance as we did not control for caffeine consumption. Caffeine is an adenosine-inhibitor that can prevent impairments in hippocampal long-term potentiation caused by sleep deprivation (Saletin et al., 2016). Thus, any effects of poor nighttime sleep may have been masked by effects of daytime caffeine consumption in our participants, particularly because MST performance has been shown to improve following caffeine intake (Borota et al., 2014). In mice, sleep-loss induced adenosine accumulation in hippocampal regions inhibited long-term potentiation, which impaired hippocampal-dependent memory (Florian et al., 2011). In our sample, adenosine accumulation in participants with shorter sleep durations across the seven days might contribute to the poorer MST performance compared to participants with longer sleep durations. However, this is only speculative.

Finally, the sample size of our exploratory study was small, which precluded examining health or demographic differences in sleep and cognition, or stratifying participants by sleep disorders. A larger sample would also allow examination of potential curvilinear relationships between sleep and performance on cognitive tests sensitive to AD (Anagnostakis et al., 2026). In addition, our sample was less ethnically diverse than the general population, and approximately 85% of participants had completed post-secondary education, which may limit the generalizability of these findings. The present study informed the design of the Brain Resilience Study, a larger investigation of resilience factors in AD that includes objective and self-reported measures of sleep alongside tests of pattern separation (McNaughton et al., 2026).

## 5. Conclusions

Our research suggests that not all cognitive tests sensitive to impairments associated with MCI and dementia may be appropriate outcome measures for clinical trials evaluating sleep-promoting treatments in older adults. In our study, the cognitive measure that was most associated with sleep quantity was the test designed to assess pattern separation performance (i.e., MST). Future studies are needed to assess whether cognitive tests designed to tax pattern separation are sensitive enough to detect cognitive improvements associated with sleep-targeting therapeutics.

## Supplementary Materials

The following supporting information can be found alongside the article.

## Author Contributions

Conceptualization, B.A.K., A.E.R. and D.G.; methodology, B.A.K., A.E.R. and D.G.; formal analysis, B.A.K., A.E.R. and D.G.; investigation, B.A.K., A.E.R. and D.G.; resources, B.A.K.; data curation, B.A.K., A.E.R. and D.G.; writing—original draft preparation, B.A.K., A.E.R. and D.G.; writing—review and editing, B.A.K. and A.E.R.; visualization, B.A.K., A.E.R. and D.G.; supervision, B.A.K.; project administration, B.A.K., A.E.R. and D.G.; funding acquisition, B.A.K. All authors have read and agreed to the published version of the manuscript.

## Institutional Review Board Statement

The research study was reviewed and approved by the Research Ethics Board at Simon Fraser University (Protocol #30000539; date of approval: 09/16/2021).

## Informed Consent Statement

The privacy rights of the participants were observed. All participants provided written informed consent to participate in the research.

## Data Availability Statement

Data is not yet available in a public repository but available upon request to the corresponding author.

## Acknowledgements

We would like to thank the following research assistants who assisted with data collection and analyses: Julia Mugliston, Kashish Mehta, Japneet Kaur, Arman Virk, Olivia Braziller, Karthikha Sri Indran, Hannah Chute, Tina Shehata, Vanessa Salzano, Sara Tam, Liliane Tansley, Robert Gibson, Alex Nash, and Miranda Chang. We would also like to thank Craig Stark (University of California, Irvine) for his invaluable advice and assistance in using the Mnemonic Similarity Task, and Ian Berkovitz (SFU) for helpful statistical consultations.

## Abbreviations

The following abbreviations are used in this manuscript:

AD: Alzheimer’s disease
MMSE: Mini Mental Status Exam
MCI: Mild cognitive impairment
MST: Mnemonic Similarity Task
DMS: Delayed Matching to Sample
CANTAB: Cambridge Neuropsychological Test Automated Battery
PVT: Psychomotor Vigilance Task
REB: Research Ethics Board
MoCA: Montreal Cognitive Assessment
ISI: Inter-stimulus interval
PAL: Paired Associates Learning
RTI: Reaction Time Inventory
SWM: Spatial Working Memory
TST: Total sleep time
IS: Interdaily stability
IV: Intradaily variability
REC: Memory Recognition
LDI: Lure Discrimination Index
L1: Lure Bin 1
L2: Lure Bin 2
RT: Reaction time
EEG: Electroencephalography

## Disclaimer/Publisher’s Note

The statements, opinions and data contained in all publications are solely those of the individual author(s) and contributor(s) and not of MDPI and/or the editor(s). MDPI and/or the editor(s) disclaim responsibility for any injury to people or property resulting from any ideas, methods, instructions or products referred to in the content.

