## Supplementary Material for "Cognitive tests of pattern separation as sleep-sensitive measures in middle-aged and older adults: An exploratory study"

**Supplementary Tables**

| **Table S1.** Sensitivity analysis of the multiple linear regression on the effect of total sleep time (TST) on MST performance | | | | | | | | | | | |
| --- | --- | --- | --- | --- | --- | --- | --- | --- | --- | --- | --- |
|  |  |  | **Unstandardized coefficients** | |  |  |  |  |  |  |  |
| **Outcome variables** | **Predictors and covariates** | **DF,error** | **B** | **Std error** | **t** |  | **F** | ***p*** | **Lower** | **Upper** | **R^2^** |
| **L1 Accuracy** |  |  |  |  |  |  |  |  |  |  |  |
| **(n=22)** |  |  |  |  |  |  |  |  |  |  |  |
|  | Constant |  | 0.51 | 0.28 | 1.87 |  |  | 0.08 | -0.06 | 1.09 |  |
|  | Age | 1,18 | -0.01 | 0.00 | -4.22 |  | 17.78 | 0.0005*** | -0.02 | -0.01 |  |
|  | Sex | 1,18 | -0.01 | 0.03 | -0.42 |  | 0.18 | 0.68 | -0.07 | 0.05 |  |
|  | TST | 1,18 | 0.08 | 0.03 | 2.82 |  | 7.97 | 0.01* | 0.02 | 0.14 |  |
|  | Model | 3,18 |  |  |  |  | 8.36 | 0.001** |  |  | 0.58 |
| **L1 False Memory Error Rate**  **(n=22)** |  |  |  |  |  |  |  |  |  |  |  |
|  | Constant |  | 0.62 | 0.28 | 2.23 |  |  | 0.04* | 0.03 | 1.21 |  |
|  | Age | 1,18 | 0.01 | 0.00 | 3.18 |  | 10.13 | 0.005** | 0.00 | 0.02 |  |
|  | Sex | 1,18 | 0.05 | 0.03 | 1.87 |  | 3.49 | 0.08 | -0.01 | 0.11 |  |
|  | TST | 1,18 | -0.08 | 0.03 | -2.96 |  | 8.77 | 0.008** | -0.14 | -0.02 |  |
|  | Model | 3,18 |  |  |  |  | 5.79 | 0.006** |  |  | 0.49 |
| Covariates: Age and sex.  Dependent variables: Lure Bin 1 Accuracy (L1 Accuracy); Lure Bin 1 False Memory Error Rate (L1 False Memory Error Rate).  Predictors: Total Sleep Time (TST).  *Note.* Type 3 partial effect tests were used. Only participants who had data from all measures were included.  * = *p* < 0.05, ** = *p* < 0.01, *** = *p* < 0.001 (two-tailed). | | | | | | | | | | | |

| **Table S2.** Multiple linear regression on the effect of total sleep time (TST) on MST performance, controlling for education | | | | | | | | | | | |
| --- | --- | --- | --- | --- | --- | --- | --- | --- | --- | --- | --- |
|  |  |  | **Unstandardized coefficients** | |  |  |  |  |  |  |  |
| **Outcome variables** | **Predictors and covariates** | **DF,error** | **B** | **Std error** | **t** |  | **F** | ***p*** | **Lower** | **Upper** | **R^2^** |
| **L1 Accuracy**  **(n=25)** |  |  |  |  |  |  |  |  |  |  |  |
|  | Constant |  | 0.37 | 0.29 | 1.28 |  |  | 0.21 | -0.23 | 0.97 |  |
|  | Age | 1,20 | -0.01 | 0.00 | -4.40 |  | 19.37 | 0.0003*** | -0.02 | -0.01 |  |
|  | Sex | 1,20 | -0.01 | 0.03 | -0.50 |  | 0.25 | 0.62 | -0.07 | 0.04 |  |
|  | Education | 1,20 | 0.00 | 0.01 | 0.58 |  | 0.34 | 0.57 | -0.01 | 0.02 |  |
|  | TST | 1,20 | 0.09 | 0.03 | 3.05 |  | 9.30 | 0.006** | 0.03 | 0.15 |  |
|  | Model | 4,20 |  |  |  |  | 7.82 | 0.0006*** |  |  | 0.61 |
| **L1 False Memory Error Rate**  **(n=25)** |  |  |  |  |  |  |  |  |  |  |  |
|  | Constant |  | 0.91 | 0.31 | 2.95 |  |  | 0.008** | 0.27 | 1.55 |  |
|  | Age | 1,20 | 0.01 | 0.00 | 2.48 |  | 6.13 | 0.02* | 0.00 | 0.02 |  |
|  | Sex | 1,20 | 0.05 | 0.03 | 1.64 |  | 2.70 | 0.12 | -0.01 | 0.11 |  |
|  | Education | 1,20 | -0.01 | 0.01 | -1.17 |  | 1.38 | 0.25 | -0.03 | 0.01 |  |
|  | TST | 1,20 | -0.08 | 0.03 | -2.59 |  | 6.73 | 0.02* | -0.14 | -0.02 |  |
|  | Model | 4,20 |  |  |  |  | 3.86 | 0.02* |  |  | 0.44 |
| Covariates: Age, sex, and education.  Dependent variables: Lure Bin 1 Accuracy (L1 Accuracy); Lure Bin 1 False Memory Error Rate (L1 False Memory Error Rate).  Predictors: Total Sleep Time (TST).  *Note.* Type 3 partial effect tests were used.  * = *p* < 0.05, ** = *p* < 0.01, *** = *p* < 0.001 (two-tailed). | | | | | | | | | | | |

| **Table S3.** Correlations between total sleep time (TST), sleep efficiency (SE), and performance on L3-L5 MST trials | | | | | |
| --- | --- | --- | --- | --- | --- |
|  | **Sleep parameters** | | | | |
| **MST outcomes** | **Average TST** | |  | **Average SE** | |
|  | *r/ρ*(df) | *p* |  | *r/ρ*(df) | *p* |
| **Participants**  **(n = 25)** |  |  |  |  |  |
| L3 Accuracy | *r(23) =* 0.21 | 0.31 |  | *r(23) =* 0.07 | 0.74 |
| L4 Accuracy | *r(23) =* 0.25 | 0.22 |  | *r(23) =* 0.06 | 0.77 |
| L5 Accuracy | *r(23) =* 0.18 | 0.38 |  | *r(23) =* 0.06 | 0.79 |
| Abbreviations: MST, Mnemonic Similarity Task; Lure Bin 3; L4, Lure Bin 4; L5, Lure Bin 5.  Correlations were run using Pearson’s correlation coefficient (*r*).  * = *p* < 0.05 (two-tailed). | | | | | |
